# Improving data archiving practices in ancient genomics

**DOI:** 10.1101/2023.05.15.540553

**Authors:** Anders Bergström

## Abstract

The sequencing of ancient DNA from preserved biological remains is producing a rich record of past genetic diversity in humans and other species. However, unless the primary data is made available in public archives in an appropriate fashion, its long-term value will not be fully realised. I surveyed publicly archived data from 42 recent ancient genomics studies. I found that half of the studies archived incomplete subsets of the generated genomic data, preventing accurate replication and representing a loss of data of potential use for future research. None of the studies met all archiving criteria that could be considered best practice. Based on these results, I make six recommendations for data producers: 1) archive all sequencing reads, not just those that can be aligned to a reference genome, 2) archive read alignments as well, but as secondary analysis files linked to the underlying raw read files, 3) provide correct experiment metadata on how samples, libraries and sequencing runs relate to each other, 4) provide informative sample metadata in the public archives, 5) publish and archive data from screening, low-coverage, poorly performing and negative experiments, and 6) document data archiving choices in papers, and review these as part of peer review processes. Given the reliance on destructive sampling of finite material, I argue that ancient genomics studies have a particularly strong responsibility to ensure the longevity and reusability of generated data.

## Introduction

The rich nature of genomic sequencing data means that new discoveries are often made using previously published data—including discoveries that the primary producers of the data might not even have envisioned. Ancient DNA has been described as a field with unusually good data sharing practices, as the vast majority of studies do share data in some form^1^. As the field has entered the era of high-throughput sequencing, however, enough attention has not always been given to how data is deposited and structured in public archives. Often, incomplete, filtered or mislabelled versions of the data are archived, meaning that published results can not be replicated from scratch by independent researchers, and that the long-term value of the data to future science is reduced.

The way in which publicly archived data is organised, accessioned and associated with useful metadata also greatly influences its utility to future users. The properties of findability, accessibility, interoperability, and reusability (FAIR) are often put forth as guiding principles for data sharing^2^. Ideally, data and associated metadata should be machine-readable and processable by automated pipelines with minimal need for manual intervention. Many users of published ancient genomics datasets will have experienced that studies in this field generally fall short of the FAIR principles, as data reuse often requires quite a bit of manual detective work, sifting through the supplementary materials of papers, and sometimes contacting the primary data producers for clarifications. Ancient genomic data will soon be available from tens of thousands of individuals, primarily from humans but also from other species as well as various environmental samples, and navigating this will become increasingly challenging if data archiving and documentation standards are not improved.

The primary public archives used for genome sequencing data are those provided by the International Nucleotide Sequence Database Collaboration (INSDC)^3^: the European Nucleotide Archive (ENA)^4^, the Sequence Read Archive (SRA)^5^, and the DNA Data Bank of Japan (DDBJ)^6^, which mirror data between each other. The INSDC ecosystem also includes the BioProject and BioSamples databases^7^. This ecosystem can be challenging to navigate, but does provide a rigorous system for the organisation and documentation of biological samples and genomic sequencing data derived from them. The discussion that follows is primarily centred around the ENA, but the principles apply more broadly.

I surveyed the genomic data deposited into public archives by 42 recent studies that sequenced ancient or historical DNA from multicellular organisms through shotgun sequencing or large-scale capture^8–49^. Studies with primarily microbial or metagenomic targets were not included here, but while current practices and considerations might differ slightly for such studies^50–52^, the fundamental archiving principles are the same whether the targeted material is genomic or metagenomic. I found widespread shortcomings in data archiving practices across the surveyed studies, and I present the results within the framework of six best-practice recommendations for producers of ancient genomic data. I also provide a condensed, step-by-step guide to archiving in line with these recommendations (**Box 1**).

## Results

### 1. Archive all sequencing reads

The most important thing to ensure is that all of the raw reads obtained in a sequencing experiment are archived, prior to any filtering or modification resulting from mapping to a reference genome, assembly, or other downstream analysis. This unfiltered version of the data represents the full information obtained from the experiment, enables any published findings to be replicated from scratch, and ensures that the long-term utility of the data to future research is maximised. The most commonly used format for raw reads is the FASTQ format^[footnote 1]^. When only reads that the data producers could successfully align to a reference genome are archived, potentially valuable data is lost from the scientific record and unavailable for reuse by future researchers.

The fundamental argument for archiving raw reads is that even if the producers of the data do not envision much additional utility for the complete set of reads, future users of the data might. Some advantages to the availability of raw reads can be recognized already:

- The ability to remap reads from scratch, e.g. with different software or parameters. Most ancient genomics studies use the bwa aln software^53^ to map reads, but if new methods that increase accuracy were to be developed, the benefit of that could only apply fully to previously published datasets if they include all reads, including those that could not be mapped by current software.
- The ability to remap reads to different reference genomes. Higher-quality reference genomes can improve read mapping accuracy and enable analyses of previously inaccessible parts of the genome. Furthermore, alternative, multiple or modified reference genomes are sometimes used in ancient genomics to overcome technical challenges^54,55^. It is likely that the process of mapping short, damaged reads to a single reference genome is a major source of reference bias in downstream analyses, as reads that display differences from the reference are more likely to end up as unmapped^56^. Methods using variation graphs, incorporating genetic variation from multiple individuals, can reduce reference bias during read mapping^57^. But if the unmapped reads from previously published studies are not available, no current or future methodological advances can reverse the reference bias introduced during the original mapping process^56,58^.
- The ability to analyse non-endogenous reads of interest, i.e. those from microorganisms and viruses. There are examples in the literature of microbial pathogen DNA being detected in previously published data^59,60^. As metagenomics detection methods improve, yet more discoveries might be made even in data that was published long ago.

Out of the 42 surveyed studies, 20 archived raw reads (**Table 1**). The remaining 22 studies only archived reads that could be aligned to a reference genome, meaning that there has been a loss of data for those studies.

**Table 1.** Survey results on the archiving of raw reads.

| Criterion | Studies meeting criterion | Studies not meeting criterion | Total number of applicable studies |
| --- | --- | --- | --- |
| Archived raw reads | 20 (47.6%) | 22 (52.4%) | 42 |
| Used capture technology, and archived raw reads | 10 (34.5%) | 19 (65.5%) | 29 |
| Trimmed adapter sequences | 9 (45%) | 11 (55%) | 20 |

Out of the 29 surveyed studies that made use of capture technology prior to sequencing, only 10 archived raw reads (**Table 1**). An argument could be made that capture technology reduces the relevance and future utility of reads not mapping to the targeted regions. However, concerns around remapping ability and reference genomes remain, and the fact that target enrichment processes are not 100% efficient means that off-target reads will be present and could prove useful. Non-targeted microorganisms have been detected in the non-endogenous reads from modern DNA capture experiments^61^. The scarcity of raw data from ancient capture experiments means that it remains largely unknown to what extent microbial DNA is detectable in such data. I argue there is no reason why raw reads from capture experiments should not be archived in the same way as raw reads from shotgun experiments.

Various processing steps are often applied to reads in ancient genomics studies: trimming off technical adaptor sequences; merging overlapping read pairs resulting from paired-end sequencing of very short molecules (and discarding those pairs that could not be merged); trimming off the last few bases of reads as these typically have higher rates of postmortem damage. No clear consensus exists on what, if any, processing to apply to reads prior to archiving. As a general principle, reads should be archived in a form that excludes technical artefacts arising from the sequencing experiment, while avoiding loss of information about the underlying biological sequence. Reads submitted for archiving should thus be demultiplexed into separate files per sample if needed, and any technical adapter sequences in the reads should be trimmed off—indeed, the ENA expects reads to not contain adapters. Out of the 20 surveyed studies that submitted raw reads, only 9 of them trimmed off adapter sequences (**Table 1**).

Further processing of the reads, however, should arguably be avoided. This includes merging of overlapping pairs in paired-end sequencing experiments. An assumption is often made that if the two reads in a pair do not overlap each other, it’s because the underlying molecule is too long to be an endogenous ancient molecule, and should thus be excluded. While this is a conservative assumption that likely benefits most downstream analyses, future users should be able to make their own choices about what reads to include or exclude. Depending on DNA preservation properties and read lengths, it should not be ruled out that some datasets might contain endogenous molecules that genuinely are longer than twice the read length and thus fail merging. Some current use cases for archived data also require the paired-end structure of reads to be intact, e.g. in metagenomics de-novo assembly^52^. In a similar vein, while analysts might choose to trim off the last few bases of all reads to mitigate the effects of postmortem damage, this should not be applied to the version of the files that are archived—it represents a loss of information about the underlying sequence, and future users should be able to make their own trimming decisions. In summary, my recommendation is thus to submit reads that have been demultiplexed and trimmed for adapters, but not merged, trimmed or filtered in any further way.

As a source of potential confusion, ENA automatically converts read alignments submitted in BAM format into unaligned reads in FASTQ format, and provides these as “*generated files*” alongside the “*submitted files*”. A user downloading an auto-generated FASTQ file might not realise that it derives from a BAM file from which unmapped or other reads have been filtered out, and thus be under the incorrect impression that it contains all the reads obtained in the experiment. Users are thus recommended to check what files were actually submitted (e.g. using the “*submitted_format*” field).

It’s worth pointing out that additional files can still be submitted to archives after a paper has been published. In principle, it is thus never too late for data producers to go back and submit missing data to existing studies, if the data still exists somewhere locally (although extra care might be needed if doing so, so as to not create additional confusion about how different files relate to each other—see recommendation 3 below).

### 2. Archive read alignments as analysis files

The standard format for representing alignments of reads to a reference genome is the BAM format^62^, along with the more heavily compressed alternative CRAM^63^. Sharing alignments should not be considered a requirement for genomics studies in the same way that sharing of raw reads is, but doing so represents a useful service. Providing the exact alignments used for published analyses, resulting from the processing and filtering deemed appropriate by the data producers, enable other researchers to quickly replicate and build upon those analyses. Archiving both raw reads and ‘analysis-level’ read alignments can thus be considered best practice in ancient genomics data sharing.

It is not ideal to simply submit read alignment BAM files to public archives alongside, and as if they were equivalent to, raw FASTQ files, however. If this is done, the BAM files will be registered as separate experiments and be assigned independent sequencing run accessions, thereby duplicating data in the archive. Instead, ENA provides an “Analysis Files” functionality which allows for read alignments to be registered as the products of secondary analysis applied to the raw reads (though these ENA analysis files are not currently mirrored to the other INSDC databases). When submitted in this way, the read alignments are linked to the corresponding raw read files, are assigned “analysis accessions” instead of run accessions, and are listed in a dedicated section on the study page.

Out of the 42 surveyed studies, 33 archived read alignments (**Table 2**). However, as discussed in the previous section, for 22 of the studies this was the only thing archived, as raw reads were not also deposited. Only 11 studies archived both raw reads and read alignments. Only 2 studies made use of the ENA “Analysis Files” functionality to appropriately categorise read alignment files as the products of secondary analysis rather than independent experiments.

**Table 2.** Survey results on the archiving of read alignments.

| Criterion | Studies meeting criterion | Studies not meeting criterion | Total number of applicable studies |
| --- | --- | --- | --- |
| Archived read alignments | 33 (78.6%) | 9 (21.4%) | 42 |
| Archived both raw reads and read alignments | 11 (26.2%) | 31 (73.8%) | 42 |
| Archived read alignments, and did so as ENA 'Analysis files' | 2 (6.1%) | 31 (93.9%) | 33 |
| Archived read alignments, and did so without any apparent hard filtering of reads | 7 (21.2%) | 26 (78.8%) | 33 |
| Archived read alignments, and did so without hard filtering on mapping quality | 21 (63.6%) | 12 (36.4%) | 33 |
| Archived read alignments, and did so without hard filtering on read length | 11 (33.3%) | 22 (66.7%) | 33 |
| Did not archive raw reads. Archived read alignments, and did so without any apparent hard filtering of reads | 5 (22.7%) | 17 (77.3%) | 22 |

The BAM format is designed to require little if any hard filtering—that is, the complete removal of reads from the file. Instead, information stored for each read on how well it is aligned to the reference, including the mapping quality metric^64^, allows for soft filtering to be performed on the fly. There is therefore little reason for the particular filtering choices preferred by the data producers to be applied to archived BAM files through hard filtering. For example, while a mapping quality threshold of 30 is often applied in ancient genomics analyses, some users might prefer to apply less stringent thresholds (for example 25^29^, 20^30^ or 10^8^). But if reads with mapping qualities below 30 have been removed from archived files, it is not possible for future users to to apply lower thresholds. Even reads with mapping qualities of 0—meaning they map equally well to more than one place in the reference genome—can be useful, for example for quantifying the copy number of a genomic region that is represented more than once in the reference. The exclusion of reads that are not uniquely mappable has impeded systematic analyses of copy number variation in ancient human genomes^65^. Reads shorter than some threshold length, typically 35 nucleotides, are similarly often excluded from analyses, but shorter reads could potentially be of future use too^66^.

Out of the 33 surveyed studies that archived read alignments in BAM format, all had removed unmapped reads from the BAM files, as noted previously^[footnote 2]^. 26 of these studies appear to have applied one or more additional hard filters (**Table 2**): 12 studies hard filtered on some mapping quality threshold of 25 or higher, and 22 studies hard filtered on some read length threshold of 30 or higher. For 22 of the 33 studies that archived read alignments, this was the only form of the data archived, as raw reads were not also archived, as noted previously (**Table 1**). Out of these, 17 applied hard filters to the BAM files, and for these studies the hard filtering of alignments thus results in the unavailability of potentially useful reads, representing additional data loss on top of that resulting from the exclusion of unmapped reads.

### 3. Provide correct experiment metadata

Ancient genomics projects often involve the sequencing of multiple libraries for the same sample, multiple sequencing runs for the same library, capture of libraries that have previously been shotgun sequenced, etc. Keeping track of how samples, libraries and sequencing runs relate to each other is important to enable correct reprocessing, and errors and ambiguity in experiment metadata is a common source of confusion for users of published ancient genomic data. A key feature of the metadata model used by the INSDC databases is that samples exist independently of studies and datasets. Samples submitted through these databases are assigned sample accessions and are also automatically registered in the NCBI BioSample database^7^. Data from multiple different experiments, even from different studies, can then be linked to a given sample accession.

Keeping track of various technical details relating to how different libraries have been constructed can also be important in downstream analyses of ancient genomics data. Different methods—e.g. commonly employed ‘double-stranded’^67^ versus ‘single-stranded’^68^ library preparation protocols, or enzymatic removal of damaged nucleotides^69^—can impact data properties in different ways. Any relevant information like this can be included in the experimental metadata when submitting data files, e.g. in the optional free text “*Library Construction Protocol*” field.

Out of the 42 surveyed studies, for 15 studies the number of sample accessions with data files linked to the study in the archive did not match the number of samples cited in the corresponding paper (**Table 3**). A user of any of these 15 datasets will thus need to engage in some detective work to figure out how archived files relate to samples described in the paper. For at least 8 studies, the discrepancy in numbers was due to experiments on the same sample, but different libraries, having been incorrectly registered as separate samples. At least 11 studies reported new data on samples for which other studies had previously reported data, but none of them associated the new data files to the existing sample accessions, but instead registered new, duplicate sample accessions. In some cases relationships to existing samples were seemingly indicated by providing sample names ending in e.g. “_new”, which does not constitute a formal link within the metadata system and is a recipe for long-term confusion. No studies indicated technical library details, such as library preparation protocol or enzymatic damage removal, in the experimental metadata submitted to the archive.

**Table 3.** Survey results on the completeness of experiment metadata accompanying archived files.

| Criterion | Studies meeting criterion | Studies not meeting criterion | Number of applicable studies |
| --- | --- | --- | --- |
| Sample number in archive matches sample number described in the paper | 27 (64.3%) | 15 (35.7%) | 42 |
| Libraries from the same sample are correctly linked to a single sample accession | 34 (81%) | 8 (19%) | 42 |
| Archived new data from previously published samples, and correctly reused existing sample accessions | 0 (0%) | 11 (100%) | 11 |
| Included technical library preparation details in archive metadata | 0 (0%) | 42 (100%) | 42 |
| BAM files have read groups | 29 (87.9%) | 4 (12.1%) | 33 |
| BAM files have SM (sample) read group fields | 27 (93.1%) | 2 (6.9%) | 29 |
| BAM files have SM read group fields, and the sample name here matches the sample name in the archive metadata | 9 (33.3%) | 18 (66.7%) | 27 |
| BAM files corresponding to a single sample contain only a single sample name in SM read group fields | 22 (81.5%) | 5 (18.5%) | 27 |
| BAM files have LB (library) read group fields | 20 (60.6%) | 13 (39.4%) | 33 |
| BAM files have LB read group fields, and the library name matches the library name in the archive metadata | 4 (20%) | 16 (80%) | 20 |
| BAM file passes validation by Picard's ValidateSamFile tool | 20 (60.6%) | 13 (39.4%) | 33 |

The importance of experiment metadata also extends to read alignments in the BAM format. The BAM format is designed to keep track of relationships between samples, libraries and sequencing runs, through the use of so-called read groups. A read group represents a set of reads that share an experimental origin: the same sample, library and sequencing run. The identity of a read group can be described in the header of the BAM file through a number of fields (“SM” for sample, “LB” for library, “PL” for sequencing platform, etc.). Most mainstream processing tools for BAM files are read group aware, and some tools will not run in the absence of certain read group fields. For example, most tools that identify PCR duplicates use the library (“LB”) read group value to only mark duplicates within a given library, even if sequenced across multiple runs, but not across different libraries from the same sample. Ideally, the sample and library IDs supplied in these read group fields within BAM files should exactly match the IDs provided in the corresponding ENA metadata, to avoid confusion and enable matching of IDs by automated processes rather than by human eyes.

Out of the 33 surveyed studies that archived BAM files, 29 made use of read groups in one way or another (**Table 3**). 27 made use of the “SM” read group field indicating sample identity of reads. However, for 18 of these studies, the sample ID supplied in the “SM” field did not match any of the sample IDs or aliases provided in the ENA metadata. For 5 studies, a single BAM file incorrectly contained more than one “SM” tag—this would, for example, cause standard variant calling software to treat different sets of reads in those files as coming from different samples. 20 of the 33 studies made use of the “LB” read group field indicating library identity of reads. However, for 16 of these studies, the library ID supplied in the “LB” field did not match the library ID provided in the ENA metadata. For 13 of the 33 studies that archived BAM files, the BAM files produced one or more errors when tested using a validation tool, which could relate to missing read groups but also other formatting issues. Overall, the archived BAM files from most of the studies would likely require manual verification or reformatting in one way or another before being safely usable in analyses.

### 4. Provide informative sample metadata

Users of publicly archived datasets also require metadata on the biological samples from which the data derives. Insufficient or incorrect metadata is a long-standing problem affecting databases on genetic biodiversity more broadly^70^. In ancient genomics studies, key bits of metadata include the geographical location where the biological remains were found, and when in the past the organism is thought to have lived (e.g. radiocarbon dates or other dating assessments). Other relevant metadata include the type of tissue that was sampled, and, for remains from human-associated archaeological sites, the name and/or cultural context of a site. Something that helps long-term data provenance is reference to any external museum, collection or excavation identifiers for the sampled specimen^71^. Many studies that do not provide particularly informative sample metadata in ENA or similar archives often do so in the publication describing the study, in which case the information is still available in the scientific record for those that look hard enough for it. However, sample metadata becomes more useful if it is also formally linked, in a standardised fashion, to the primary data in the public archives.

Out of the 42 surveyed studies, 25 provided a geographical label of one kind or another in the sample metadata submitted to public archives, but only 12 used a dedicated metadata field for this (typically “*geographic location (region and locality)*”) (**Table 4**). Only 3 studies provided latitude and longitude coordinates. 7 studies provided some kind of dating information, but only 1 used a dedicated metadata field for this (“*date*”), and only 1 provided the radiocarbon lab codes that serve as unique identifiers for radiocarbon dates. 15 studies provided some information on what tissue was sampled (though in 3 cases this was not higher resolution than just “Bone” versus “Tooth”), but only 10 studies provided tissue information in a dedicated metadata field (typically “*tissue*”), while the others provided it in the general purpose “*Description*” field. 6 out of the 42 studies provided some kind of information on the identity or source of the sampled specimen (using the “*biological material*” or “*specimen voucher*” fields), though identifiers were not always accompanied by information on what external collection or museum those identifiers refer to.

**Table 4.** Survey results on the completeness of sample metadata accompanying archived files.

| Criterion | Studies meeting criterion | Studies not meeting criterion | Number of applicable studies |
| --- | --- | --- | --- |
| Metadata contains any geographical information | 25 (59.5%) | 17 (40.5%) | 42 |
| Metadata contains geographical information, and represents this in a dedicated metadata field | 12 (48%) | 13 (52%) | 25 |
| Metadata contains geographical information, and includes geographical coordinates | 3 (12%) | 22 (88%) | 25 |
| Metadata contains any information on dating of the sample | 7 (16.7%) | 35 (83.3%) | 42 |
| Metadata contains dating information, and represents this in a dedicated metadata field | 1 (14.3%) | 6 (85.7%) | 7 |
| Metadata contains dating information, and includes radiocarbon lab codes | 1 (14.3%) | 6 (85.7%) | 7 |
| Metadata contains any information on sampled tissue | 15 (35.7%) | 27 (64.3%) | 42 |
| Metadata contains tissue information, and represents this in a dedicated metadata field | 10 (66.7%) | 5 (33.3%) | 15 |
| Metadata contains any reference to the museum/collection/excavation origin or identity of the sample | 6 (14.3%) | 36 (85.7%) | 42 |

There is currently no clear consensus in ancient genomics on what metadata should be reported, or in what format various types of metadata should be provided. There is a strong need for standardisation in this area, perhaps including the adoption of controlled vocabularies (e.g. for tissues^72^), but that is a discussion that is outside the scope of this article. Ongoing efforts to e.g. establish a MIxS (*Minimum Information about any (x) Sequence*^*73*^) extension checklist for ancient samples would be very valuable in this regard^74^. One reason for the poor state of sample metadata in the public archives is arguably that the standard metadata fields in the INSDC databases are not geared towards ancient samples, such that it’s not necessarily clear to sample submitters in what fields different pieces of information should go. Once stable metadata standards have emerged in the ancient genomics community, these could likely also be adopted by the ENA through the creation of a dedicated sample checklist (the ENA has already adopted such checklists for various other types of samples with particular metadata requirements). In the meantime, a minimal recommendation to data submitters is: whatever metadata you include in the supplementary materials of the paper, make an effort to submit that same exact metadata to the public archive too, use metadata fields that are as appropriate as possible, and strive to make information as machine-readable as possible.

It’s worth pointing out that, as samples exist independently of studies and data files, sample metadata in the INSDC databases can be updated even after data files have been submitted. It is thus never too late to go back to previously registered samples and supplement them with more informative metadata. Lastly, from mid-2023 the INSDC databases have made some minimal spatiotemporal metadata mandatory for newly submitted samples^75^. However, it should be noted that this minimal metadata concerns when and where (e.g. a museum) a sample was collected with the intention of DNA sequencing, and not the time and place where the organism lived^76^. This might become a source of confusion for ancient samples, and additional metadata fields will thus still be needed to provide this crucial geographical information for ancient samples.

### 5. Archive data from low-coverage, screening and negative experiments

Many biological remains do not contain enough endogenous DNA to make genome sequencing worthwhile. Many ancient genomics projects first screen samples though very low-coverage sequencing, before deciding on what samples to perform deeper sequencing on. Data from this screening phase is typically not used for published analyses, and often not deposited in public archives. Some papers mention how many samples were screened but not taken forward, but this is not always done. In a similar way, studies that do not necessarily employ a screening phase will sometimes nonetheless choose to exclude some samples from a publication because the data obtained from them was poor, and not archive the data from those experiments. While difficult to estimate, it is a possibility that data from the majority of ancient genome sequencing experiments performed to date actually remain unpublished and unavailable.

It could be argued that if one undertakes destructive sampling of any ancient biological remains, doing justice to those remains should involve a commitment to publish and archive the resulting data, whatever its quality turns out to be^[footnote 3]^. The practice of not archiving screening and other low-coverage or poor data can be seen as another instance of when data producers do not envision any utility of the data, but future users might. Already, several potential uses for this kind of sequencing data from ancient remains can be identified:

- Microbial and viral DNA can sometimes be detectable even in very low-coverage data. Detection in screening data often informs decisions on capture experiments targeting pathogenic microbes of interest. Beyond these, the non-pathogenic microbial content and what it says about post-mortem degradation processes remains largely unexplored. With continued advances in computational metagenomics methodology, and increased understanding of past pathogenic and non-pathogenic microbial diversity, samples that were initially deemed uninteresting might well be found to contain data of interest by future researchers.
- Data from samples with little or no endogenous DNA can contribute to the understanding of factors influencing DNA preservation. If data from large numbers of such ‘negative’ samples and their metadata was available, supervised models could be trained to predict DNA preservation as a function of various sample properties. Currently, however, the published literature primarily just contains results from relatively successful samples, thereby hindering systematic explorations in this area.
- Determination of biological sex through coverage analyses of the sex chromosomes does not require very high sequencing coverage, and is sometimes possible even for relatively poorly preserved samples^77,78^. While not necessarily very useful in most genomics analyses, the biological sex of individuals across regions, periods, sites etc. can inform on social, behavioural and cultural aspects^79–82^.

Out of the 42 surveyed studies, 32 mentioned in the paper the total number of studied samples but did not necessarily archive data from all samples (**Table 5**). 19 of the studies appear to have archived data from all studied samples, even if some were then left out of downstream analyses. For the remaining 23 studies, data from samples other than those included in the published analyses, whether such samples were explicitly mentioned or not discussed at all, did not seem to have been archived—meaning that data from some of the experiments conducted as part of these studies likely remain unpublished and unavailable^[footnote 4]^.

**Table 5.** Survey results on the reporting and archiving of negative or poorly performing experiments.

| Criterion | Studies meeting criterion | Studies not meeting criterion | Number of applicable studies |
| --- | --- | --- | --- |
| Paper mentions the total number of studied samples, even if some were left out from analyses | 32 (76.2%) | 10 (23.8%) | 42 |
| Data from all studied samples appears to have been archived, even if some were left out from analyses | 19 (45.2%) | 23 (54.8%) | 42 |

Beyond questions of reproducibility and future utility, the practice of only publishing results on ‘positive’ and not ‘negative’ samples also has implications for the interpretation of scientific results. This is especially true for studies reporting the detection of pathogen DNA in ancient remains. If results from negative samples are not reported, the prevalence of a given pathogen across time and space can not be assessed. Some studies do mention the total number of screened samples, e.g. a study on hepatitis B described data from 137 samples produced after initially screening 7918 human samples of unspecified origin^83^, and a study on smallpox described data from 13 samples produced after initially screening 1867 human samples from Eurasia and the Americas from the last ∼32,000 years^84^. The latter study also reported results on the number of positive and negative samples across broad time periods, thus providing valuable evidence on past prevalence. It could be argued that published results should not refer to data that is not publicly archived, as other researchers can not reproduce those results. I would instead argue that transparency about what was screened is never a bad thing, especially if it helps interpretation of the scientific findings. Even more useful, of course, would be if all the screening data is published and archived.

Some highly iconic and important human and other biological remains have presumably been sampled for DNA without success, but those negative results have not been described in the scientific literature. It would arguably be valuable to make the apparent lack of endogenous DNA in some of these high-profile specimens part of the public, scientific record, and to make the screening data that was nonetheless produced available for anyone who wants to analyse its properties independently.

### 6. Document and review archiving choices

Many of the common archiving shortcomings discussed here could be overcome relatively easily. A natural mechanism for the flagging of issues relating to the public archiving of data would be if editors and reviewers of papers scrutinised this aspect too, alongside their review of the scientific results of a paper, but this is rarely done. Data availability statements and methods sections typically contain little or no information on archiving other than just the archive accession number. Providing more details on what files were actually archived, whether any filters were applied etc., would both help future users and allow reviewers to scrutinise the archiving choices. Papers submitted for review often withhold the archive study accession number until the paper is accepted, or authors set the study itself to not be publicly visible until the paper is published. Some authors might not be comfortable with making their data publicly available pre-publication, but doing so for the review process would also contribute towards better standards in the field. It’s also worth noting that this does not need to entail making any findings public pre-publication—the study description texts in ENA/SRA can be updated over time, and do not need to correspond to paper abstracts, or necessarily mention any scientific findings from the study at all. In fact, an archive study description is arguably more useful if it focuses on the technical details of the archived data, rather than simply duplicating the abstract of a published paper.

## Conclusions

Shortcomings in how ancient genomics data is archived is likely primarily the result of data submitters not being very familiar with how the public archives are structured, and the submission processes being quite complex. Some studies do put substantial efforts into providing comprehensive information on samples, libraries and sequencing experiments in the supplementary materials of papers, or in custom files hosted on institutional or other websites. However, a reliance on inconsistently structured supplementary materials, non-permanent websites, and direct requests to authors, is not a rigorous long-term solution for a data-intensive research field. Rather than treating the task of data archiving as a necessary inconvenience to meet a journal requirement, I would urge researchers in ancient genomics to treat this component of their research projects with the same level of attention and rigour as they treat other components. Building the steps necessary for appropriate archiving into standardised pipelines and workflows could be one way to make this process easier.

During the Human Genome Project, the so-called Bermuda principles prescribed the pre-publication release of genomic data as soon as it was produced. Along with the Fort Lauderdale Agreement that aimed to prevent data producers being scooped on principal findings, this has nurtured a long-standing culture of open and early data sharing in genomics^85^. Upholding these norms, and making appropriate use of existing archive infrastructure, will be increasingly important as ancient genomics continues to grow rapidly. The dependence on destructive sampling of finite and sometimes very precious material arguably means that ancient genomics studies have a particularly strong responsibility to ensure that data is appropriately archived and reusable.

### Box 1

**Step-by-step best practices for archiving ancient genomics data in the European Nucleotide Archive (ENA)**

1. Register samples, to obtain sample accession numbers. Register and archive data even for samples in your study that performed poorly in terms of endogenous DNA recovery and could not be analysed further. If other data from a given sample has already been archived, e.g. by a previous publication, identify and reuse the existing sample accession number rather than registering a new accession number.
2. Provide as much useful metadata on the registered samples as possible, including on: geographical origin (including geographical coordinates), dating information (including radiocarbon lab codes if available), cultural context if relevant, tissue type, and any external references to museum or collection identifiers of specimens.
3. Prepare raw reads (preferably in FASTQ format, though can also be BAM/CRAM). Demultiplex reads, trim off residual adaptor sequences, but ideally do not merge overlapping read pairs from paired-end experiments. Do not filter or trim reads on length or base quality properties.
4. Deposit raw reads. Each FASTQ file submitted to the archive should correspond to a set of reads that come from the same sample, library and sequencing run. Do not concatenate reads that come from the same sample but from different runs or libraries into a single FASTQ file. Instead, deposit separate files containing data from the same sample separately and link them to the same, single sample accession number.
5. Provide as much useful experimental metadata on the FASTQ files as possible, including on: library preparation strategy (e.g. double-stranded vs single-stranded), any enzymatic damage removal steps performed, any size selection steps performed.
6. Prepare read alignments, if your study used them (BAM/CRAM format). Do not exclude reads from BAM files on the basis of mapping quality, read length, duplicate status or other properties. Only exclude unmapped reads if raw read files have already been archived, otherwise include unmapped reads in the BAM files too. Only recalibrate or otherwise modify base qualities in the BAM file if raw read files have already been archived. Make sure the BAM files have read groups that accurately indicate the origin of each read (sequencing run, library ID, sample ID, sequencing platform). Make sure the identifiers used in the read groups inside the BAM files match the identifiers used in the metadata for the corresponding raw read (FASTQ) files. Run a BAM validation tool (e.g. Picard’s ValidateSamFile) to identify any formatting issues.
7. Deposit read alignments. Deposit one BAM/CRAM file per sample, including reads from multiple sequencing runs and/or libraries if relevant. If raw reads were already deposited, then deposit read alignments as “Analysis Files”, not as primary data files, and link them to the corresponding raw read files. If unaligned reads were not deposited separately, then deposit the BAM/CRAM files as the primary data files.
8. Describe in detail what data was archived in the data availability statement of the paper (or if there is no such section, in the methods section). Describe if data was archived for all samples described in the paper, or only the subset that were carried fourth for the final analyses. Describe if raw reads, read alignments, or both were archived. Describe if any filtering was performed on the archived files.

## Methods

Studies to include in the data survey were identified as follows: I looked for papers published in 2021, 2022 and 2023 in the journals Nature, Science and Cell, by searching for “ancient DNA” or “ancient genome” using Google Scholar and the journals’ own search tools. I included studies reporting shotgun or genome-wide capture data (but not e.g. only mitochondrial capture data) targeting any multicellular organism, but not studies focussed on microbial or metagenomic targets. This identified 42 studies, but it is possible that I missed some. I downloaded experimental metadata on the archived data files for each study from the ENA (www.ebi.ac.uk/ena/, accessed February 15th, 2023 for studies published in 2021-2022, and accessed January 16th, 2024 for studies published in 2023). I obtained associated metadata from the NCBI BioSample database, downloading the full metadata from all BioSample samples linked to the given ENA study / BioProject accession (https://www.ncbi.nlm.nih.gov/biosample/, accessed February 15th, 2023 for studies published in 2021-2022, and accessed January 16th, 2024 for studies published in 2023). Three studies had archived data in the GSA database^86^ rather than in the ENA/SRA, and I obtained corresponding information from there (https://ngdc.cncb.ac.cn/gsa/, accessed February 15th, 2023). To survey properties of read and read alignment files, I downloaded the first FASTQ and/or the first BAM file listed in the ENA/GSA for each study. If there are differences in properties between files within the same study, this will thus not have been picked up here. Two studies had deposited raw, unaligned reads in BAM format rather than FASTQ format, and I treated these as equivalent to FASTQ files for reporting purposes. Information was extracted from BAM files using samtools^62^. BAM validation was performed using Picard’s ValidateSamFile^87^.

## Data availability

The data analysed in this study was obtained from the following archive study accessions: PRJEB54831 (ENA)^8^, PRJEB51180 (ENA)^9^, PRJCA005576 (GSA)^10^, PRJEB52849 (ENA)^11^, PRJEB42656 (ENA)^12^, PRJEB52230 (ENA)^13^, PRJNA786530 (SRA)^14^, PRJNA1005336 (SRA)^15^, PRJEB51440 (ENA)^16^, PRJEB56773 (ENA)^17^, PRJEB49291 (ENA)^18^, PRJEB42781 (ENA)^19^, PRJEB46734 (ENA)^20^, PRJNA798381 (SRA)^21^, PRJEB42269 (ENA)^22^, PRJEB47891 (ENA)^23^, PRJEB54899 (ENA)^24^, PRJEB43715 (ENA)^25^, PRJEB39134 (ENA)^26^, PRJEB46162 (ENA)^27^, PRJEB46875 (ENA)^28^, PRJEB44430 (ENA)^29^, PRJEB42199 (ENA)^30^, PRJEB38555 (ENA)^31^, PRJEB55327 (ENA)^32^, PRJEB56213 (ENA)^33^, PRJEB51862 (ENA)^34^, PRJEB58698 (ENA)^35^, PRJEB62503 (ENA)^36^, PRJEB66319 (ENA)^37^, PRJEB59008 (ENA)^38^, PRJEB61818 (ENA)^39^, PRJEB50368 (ENA)^40^, PRJEB50857 (ENA)^41^, PRJCA003870 (GSA)^42^, PRJCA003699 (GSA)^43^, PRJEB53475 (ENA)^44^, PRJEB37782 (ENA)^45^, PRJNA687817 (SRA)^46^, PRJEB42372 (ENA)^47^, PRJEB66422 (ENA)^48^, PRJEB57364 (ENA)^49^.

## Acknowledgements

The author thanks James Fellows Yates, Josephine Burgin, Torsten Günther, Thiseas Lamnidis, Kendra Sirak and Pontus Skoglund for constructive comments on the manuscript.

## Footnotes

1 The BAM format can also store unmapped reads, such that an unfiltered BAM file can contain the same complete information as a FASTQ file. If unfiltered BAM files that contain unmapped reads are archived, as is often done in modern genomics studies, it would not be necessary to also archive FASTQ files. Indeed, the ENA’s primary recommendation is to deposit reads in the BAM or CRAM formats. However, I would recommend that ancient genomics studies deposit FASTQ files, as they are directly usable by most common software and reduce ambiguity regarding whether all raw reads are in fact present.

2 An argument for hard filtering of BAM files is that it reduces file sizes, but I would argue that such gains are typically modest and do not justify the long-term data loss. The one hard filtering step that often does lead to substantial file size reductions is the removal of unmapped reads, owing to the large amounts of non-endogenous DNA typically recovered in ancient sequencing data. If raw reads are not also submitted separately, as discussed above, removing unmapped reads entails a loss of data and should thus be avoided. But if raw reads are also submitted separately as recommended here, there is no loss of data, and in that case providing smaller, analysis-ready BAMs without unmapped reads is arguably the most practical for users.

3 Another aspect of relevance is authorship fairness—archaeologists, museum curators etc. that provide samples that turn out negative for DNA will often not end up as authors on the publications that emerge from projects, even if their contributions were the same as those of providers of samples that turned out positive for DNA.

4 There are practical challenges to sharing screening and negative data. For some samples there might not be a suitable paper where they would naturally fit in, especially if all samples from a given project or archaeological site were negative. Even in such cases, data could be published in stand-alone publications, e.g. in archaeology journals as reports on DNA preservation at a set of sites, or in journals offering data-focussed articles that do not require interpretation of results (e.g. GigaScience, Scientific Data, F1000 Research). Stand-alone ENA study accessions could also be used to archive screening data, and these would not necessarily have to be linked to single publications—for example, a laboratory or project could create an ENA study titled “Ancient genome screening data from laboratory X”, simply deposit all of their screening data there, and provide the study accession number in multiple publications.

## Notes

### Competing Interest Statement

The authors have declared no competing interest.

### Summary of Updates

Data survey expanded to cover papers published in 2023; Box 1 with step-by-step guide added

